# Frequency- and state-dependent effects of hippocampal neural disinhibition on hippocampal local field potential oscillations in anaesthetised rats

**DOI:** 10.1101/2019.12.18.880864

**Authors:** Miriam Gwilt, Markus Bauer, Tobias Bast

## Abstract

Reduced inhibitory GABA function, so-called neural disinhibition, has been implicated in cognitive disorders, including schizophrenia and age-related cognitive decline. We previously showed in rats that hippocampal disinhibition by local microinfusion of the GABA-A antagonist picrotoxin disrupted memory and attention and enhanced hippocampal multi-unit burst firing recorded around the infusion site under isoflurane anaesthesia. Here, we analysed the hippocampal LFP recorded alongside the multi-unit data. We predicted frequency-specific LFP changes, based on previous studies implicating GABA in hippocampal oscillations, with the weight of evidence suggesting that disinhibition would facilitate theta and disrupt gamma oscillations. Using a new semi-automated method based on the kurtosis of the LFP peak-amplitude distribution as well as on amplitude envelope thresholding, we separated three distinct hippocampal LFP states under isoflurane anaesthesia: ‘burst’ and ‘suppression’ states – high-amplitude LFP-spike bursts and the interspersed low-amplitude periods – and a medium-amplitude ‘continuous’ state. The burst state showed greater overall power than suppression and continuous states and higher relative delta/theta power, but lower relative beta/gamma power. The burst state also showed reduced functional connectivity across the hippocampal recording area, especially around theta and beta frequencies. Overall neuronal firing was higher in the burst than the other two states, whereas the proportion of burst firing was higher in burst and continuous states than the suppression state. Disinhibition caused state- and frequency-dependent LFP changes, tending to increase power at lower frequencies (<20Hz), but to decrease power and connectivity at higher frequencies (>20Hz) in burst and suppression states. The disinhibition-induced enhancement of multi-unit bursting was also state-dependent, tending to be more pronounced in burst and suppression states than the continuous state. Overall, we characterized three distinct hippocampal LFP states in isoflurane-anaesthetized rats. Disinhibition changed hippocampal LFP oscillations in a state- and frequency-dependent way. Moreover, the disinhibition-induced enhancement of multi-unit bursting was also LFP-state dependent.

## 1. INTRODUCTION

Subconvulsive neural disinhibition, i.e. reduced inhibitory GABA function, within the hippocampus has been implicated in important neuropsychiatric disorders, including schizophrenia and age-related cognitive decline, and in the cognitive impairments characterizing these disorders (Bast, Pezze, & McGarrity, 2017; Benes & Berretta, 2001; Heckers & Konradi, 2015; McGarrity, Mason, Fone, Pezze, & Bast, 2017; Nava-Mesa, Jiménez-Díaz, Yajeya, & Navarro-Lopez, 2014; Palop & Mucke, 2016; Stanley, Fadel, & Mott, 2012; Thomé, Gray, Erickson, Lipa, & Barnes, 2016). In a recent study (McGarrity et al., 2017), we pharmacologically disinhibited the temporal (also known as ventral) to intermediate hippocampus in rats by acute local microinfusion of the GABA-A antagonist picrotoxin. Such hippocampal disinhibition caused clinically-relevant cognitive impairments, including in hippocampal memory function and in attentional performance that relies on prefrontal-striatal mechanisms, consistent with the idea that hippocampal disinhibition disrupts both hippocampal processing and processing in hippocampal projection sites (Bast et al., 2017). In addition, electrophysiological recordings around the infusion site under isoflurane anaesthesia showed that disinhibition enhanced burst firing of hippocampal neurons, as reflected by multi-unit data. The purpose of the present paper is to report the analysis of the impact of disinhibition on the hippocampal local field potential (LFP), which we recorded alongside the multi-unit data.

Brain rhythms or oscillations, i.e. synchronized changes in the activity of many neurons, as revealed by LFP recordings have been suggested to be important for cognitive processing, including memory, because they bind neurons into functional assemblies (Buzsáki & Draguhn, 2004; Colgin, 2016). Alterations in brain rhythms have been reported in many neuropsychiatric disorders and have been suggested to arise partly from GABA dysfunction (Uhlhaas & Singer, 2006, 2010). Two prominent, widely studied hippocampal LFP rhythms are the theta and gamma rhythms. In the rat hippocampus, theta ranges from 4-12 Hz, whereas gamma oscillations range from 25-100Hz, including ‘slow’ or ‘low’ gamma from 25-55 Hz (also known as beta rhythms) and ‘fast’ or ‘high’ gamma’ from 60-100 Hz (Colgin, 2016). Substantial evidence suggests that theta and gamma LFP rhythms depend on hippocampal GABAergic inhibition, which will be considered in the following two paragraphs. Another prominent hippocampal rhythm, sharp-wave ripples (110–250 Hz ripples superimposed on 0.01–3 Hz sharp waves) (Colgin, 2016), will not be considered further in this paper, because it is not expressed under volatile anaesthetics (Ylinen et al., 1995) and the frequency range of our LFP recordings (0.7-170Hz) did not encompass the full range of sharp waves and ripples.

As to hippocampal theta, septal projections to the hippocampus, including excitatory cholinergic projections and GABAergic projections, which disinhibit the hippocampus by innervating hippocampal inhibitory GABA interneurons (Borhegyi, Varga, Szilagyi, Fabo, & Freund, 2004; Freund & Antal, 1988; Hangya, Borhegyi, Szilágyi, Freund, & Varga, 2009; Tóth, Freund, & Miles, 1997), are thought to be a main driver (Colgin, 2016). In line with this, septal inactivation reliably abolishes hippocampal theta in freely moving rats (Brandon et al., 2011; Koenig, Linder, Leutgeb, & Leutgeb, 2011) and in urethane-anaesthetized rats with septal inactivation, hippocampal theta could be reinstated by cholinergic stimulation or disinhibition (with the GABA-A antagonist bicuculline) of the hippocampus (Smythe, Colom, & Bland, 1992). Moreover, cholinergic stimulation of rat hippocampal slices by carbachol reliably induces theta in vitro, and pharmacological disinhibition by GABA-A antagonists has been reported to enhance the amplitude of such cholinergically induced theta oscillations (Golebiewski, Eckersdorf, & Konopacki, 1996; Konopacki, Gołebiewski, Eckersdorf, Błaszczyk, & Grabowski, 1997; Kowalczyk, Bocian, & Konopacki, 2013). In line with such theta-enhancing effects of hippocampal disinhibition, hippocampal inhibition with the GABA-A agonist muscimol disrupted cholinergically induced theta in the rat hippocampus in vitro (Golebiewski et al., 1996) and in vivo (Smythe et al., 1992). These findings thus support an inverse relationship between hippocampal GABA function and theta amplitude (Kowalczyk et al., 2013).

However, there is also evidence that some types and properties of hippocampal theta are positively modulated by GABA function and disrupted by hippocampal disinhibition. Genetic ablation of synaptic inhibition of parvalbumin-positive GABAergic interneurons disrupted hippocampal theta in freely moving mice (Wulff et al., 2009). GABA-A antagonists reduced the power of theta induced by septal activation in an in vitro septo-hippocampal preparation from rats (Goutagny, Manseau, Jackson, Danik, & Williams, 2008), and of nicotine-induced (Lu & Henderson, 2010) and electrically induced (Heynen, Sainsbury, & Bilkey, 1993) theta in rat hippocampal slices, and decreased frequency and coherence (connectivity) of carbachol-induced theta between the entorhinal cortex and subiculum in vitro (Levesque, Cataldi, Chen, Hamidi, & Avoli, 2017). Moreover, optogenetic activation of parvalbumin-positive GABAergic hippocampal interneurons strengthened, whereas silencing of these neurons disrupted, theta in an in vitro preparation of the mouse hippocampus with intact intrinsic but severed extrinsic connectivity, suggesting that GABAergic inhibition by parvalbumin neurons supports intrinsically generated theta (Amilhon et al., 2015). Finally, genetic knock-out of the GABA-A receptor subunit β_3_ decreased power, frequency and regularity of hippocampal theta and theta cross-correlation between different hippocampal subfields in freely moving mice (Hentschke et al., 2009).

There is also substantial evidence linking hippocampal gamma oscillations to inhibitory GABAergic hippocampal interneurons (Colgin, 2016). In rodents, firing of hippocampal interneurons is phase-locked to spontaneous hippocampal gamma oscillations (Belluscio, Mizuseki, Schmidt, Kempter, & Buzsaki, 2012; Mann, Suckling, Hajos, Greenfield, & Paulsen, 2005; Tiesinga, Fellous, José, & Sejnowski, 2001; Traub et al., 2000; Tukker, Fuentealba, Hartwich, Somogyi, & Klausberger, 2007), and intracellularly recorded inhibitory postsynaptic potentials of pyramidal cells, recorded in vivo, are reflected in gamma oscillations (Penttonen, Kamondi, Acsady, & Buzsaki, 1998; Soltesz & Deschenes, 1993). This indicates that hippocampal gamma is generated by hippocampal GABA interneurons (Colgin, 2016). Consistent with this, GABA-A antagonists have been shown to disrupt different types of experimentally induced gamma oscillations in hippocampal slices, with the magnitude of this effect depending on the hippocampal subfield (Bartos, Vida, & Jonas, 2007; Fisahn et al., 2004; Traub, Bibbig, LeBeau, Buhl, & Whittington, 2004; Whittington, Traub, Kopell, Ermentrout, & Buhl, 2000).

Overall, the reviewed evidence on the GABAergic mechanisms of hippocampal theta and gamma oscillations suggests that pharmacological hippocampal disinhibition may alter both oscillations, with the weight of evidence suggesting that gamma amplitude may be reduced, whereas theta amplitude may be enhanced. However, there is also evidence suggesting that some features of the theta rhythm, including its coherence across the hippocampus, may be disrupted.

**Figure 1:**
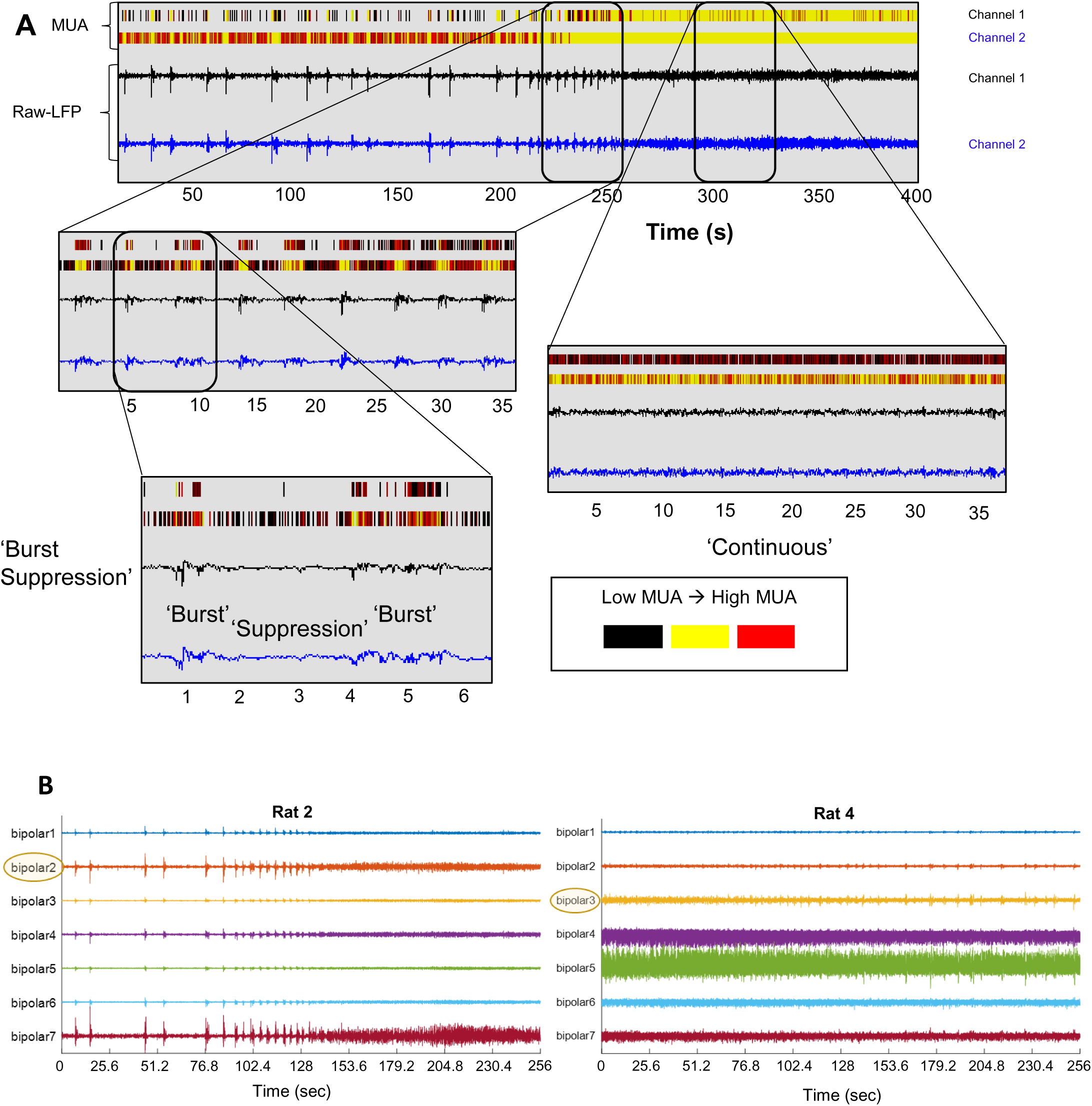
Three distinct LFP states in the rat hippocampus under isoflurane anaesthesia. **A** The top panel shows two 400-s long LFP traces, with corresponding multi-unit activity (MUA), recorded simultaneously from the temporal to intermediate hippocampus (using an 8-wire electrode array) under isoflurane anaesthesia. They illustrate the transition from the burst-suppression state (left), comprising burst and suppression states, to the continuous state (right). The middle panel shows the 40-s sections indicated in the top panel with an expanded time line. The bottom panel depicts a 7-s period of the burst-suppression state with an expanded time line, showing the LFP signal during burst and suppression states in higher detail. The three distinct LFP states also differ with respect to multi-unit spiking, particularly MUA tends to be higher during the burst than in the suppression state. **B** The two plots show the LFP traces recorded at the 7 bipolar channels in two different rats. Both examples show the burst and suppression states, as well as the continuous state: in the example shown on the top, burst-suppression precedes the continuous state, whereas in the bottom the continuous state precedes burst-suppression. In the example recordings shown in the top, the three distinct LFP states are visually more clearly separated than in the example shown in the bottom. The example traces shown on the top include those shown in Figures 1A, from rat number 2, where the states are visually clearly separated; bipolar channel 2 was selected as the ‘LFP state-defining’ channel because of very clear visual separation of the three states based on the amplitude differences. In the example LFP traces shown in the bottom, from rat number 4, the three LFP states are visually less clearly separated. Channel number 3 was selected as the ‘LFP state-defining’ channel for this rat because, although it has smaller overall amplitudes than other channels, the three different states are visually more clearly separated, especially compared to channels 4-7.

When analysing LFP recordings from anaesthetized rodents, it is important to consider that, under anaesthesia, many brain regions, including the hippocampus, show different LFP states characterized by distinct LFP patterns which can alternate (Clement et al., 2008; Kenny, Westover, Ching, Brown, & Solt, 2014; Land, Engler, Kral, & Engel, 2012; Lustig, Wang, & Pastalkova, 2016; Wolansky, Clement, Peters, Palczak, & Dickson, 2006) (for an example of our hippocampal recordings, see **Figure 1A**). One key LFP state observed with many anaesthetics, including isoflurane, is the burst-suppression state. The burst–suppression state refers to periods of LFP ‘bursts’ with high amplitude deflections interspersed between quiet periods of low amplitude LFP signal (= ‘suppression’ phases). Another type of LFP state is characterised by a more continuous LFP pattern with continuous LFP activity of lower amplitude than during LFP ‘bursts’. The balance between the burst-suppression and continuous states changes with anaesthetic depth, with burst-suppression tending to become more dominant the deeper the anaesthesia (Land et al., 2012; Lustig et al., 2016). To analyse the impact of neuropharmacological manipulations, including hippocampal disinhibition, on LFP oscillations under anaesthesia it is important to separate these different LFP states for two main reasons. First, the large-amplitude LFP signals characterizing the burst state reflect major non-stationarities in the otherwise low-amplitude signals and may thus dominate the power spectral analysis, occluding frequency components that may be more characteristic of the other LFP states. Second, given the distinct characteristics of LFP patterns in different LFP states, neuropharmacological mechanisms underlying oscillatory activity may be LFP-state dependent.

In the present study, we aimed to characterise the effect of hippocampal disinhibition, by picrotoxin infusions into the temporal to intermediate hippocampus, on hippocampal neural oscillations around the infusion site using the LFP data recorded by McGarrity et al. (2017). First, we developed an objective, semi-automated method to separate the three distinct LFP states described above (the continuous LFP state from the burst-suppression state, and, within the latter, the burst from the suppression state; **Figure 1A**) and compared several properties of these states (frequency-specific power and connectivity, and multi-unit parameters). Second, we examined the impact of picrotoxin on hippocampal LFP properties (frequency-specific power and connectivity) in the three distinct LFP states. Third, we examined if the disinhibition-induced enhancement of multi-unit burst firing reported previously (McGarrity et al., 2017) is dependent on the LFP state.

## 2. METHODS

### 2.1 LFP and multi-unit data recording and pre-processing

Analysis was carried out on the hippocampal LFP and multi-unit data collected in our previous study (McGarrity et al. 2017). In brief, LFP and multi-unit data were recorded simultaneously in isoflurane anesthetized male adult (2-3 months) Lister-hooded rats, using a custom-made assembly of a 33-gauge stainless steel infusion cannula and an 8-electrode (microwire) recording array that was implanted into the hippocampus (**Figure 2**, top left). The cannula tip, through which saline or picrotoxin solution could be injected, touched the electrodes and was positioned about 0.5 mm above the tips of the central electrodes. The assembly was stereotactically implanted into the right hippocampus, such that the 8-electrode array was arranged perpendicular to the brain midline and anterior to the infusion cannula, with the cannula tip aimed at coordinates in the right temporal (also known as ventral) to intermediate hippocampus (5.2 mm posterior to bregma, 4.8 mm lateral from midline, and 6.5 mm ventral from dura).The 8-electrode array spread approximately 2 mm, in the medio-lateral direction (**Figure 2**, top right). The extracellular signal recorded by the electrodes was band-pass filtered into LFP (0.7 and 170 Hz) and multi-unit (250 Hz to 8kHz) data, which were recorded for a 30-min baseline period and a 60-min period following infusion (over about 1 min) of either 0.5ul saline (n = 7 rats) or 150ng/0.5ul picrotoxin (n = 6 rats) in a between-subjects design. All data were collected continuously and saved in 5 minutes bins, i.e. six pre-infusion (baseline) and 12 post-infusion 5-min bins. Although the picrotoxin group in the original study included eight rats (McGarrity et al. 2017), only six of these rats could be included in the present analysis, because one of the datasets could not be read into MATLAB for analysis and another dataset was recorded using a bundle array and, therefore, could not be combined with the rest of the data for LFP analysis.

**Figure 2:**
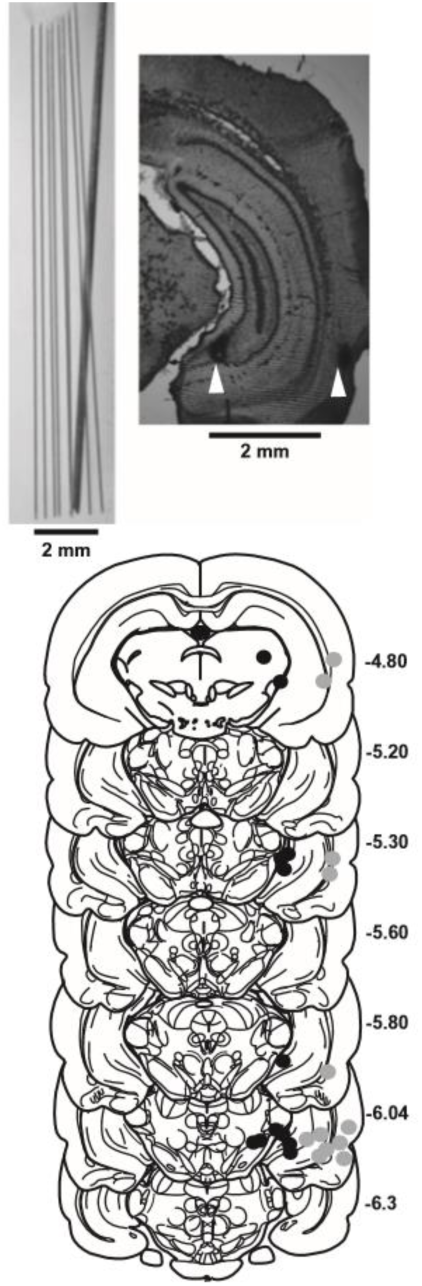
Placement of the infusion-recording array within the temporal to intermediate hippocampus. The top pictures show, left, the 8-electrode array with the attached infusion cannula and, right, an example coronal section through the hippocampus, with the placements of the most medial and most lateral electrodes indicated by the white arrow heads. The array was arranged perpendicular to the midline of the brain, with the infusion cannula located just posterior to the center of the array. In the bottom, the most medial (black dots) and most lateral (grey dots) electrode placements are indicated for all rats included in the analysis on drawings of coronal brain sections taken from the atlas by (Paxinos & Watson, 1998). Note that the most medial and/or the most lateral electrodes of the 2-mm 8-electrode recording array were located outside of the hippocampus (typically 1–3 electrodes per rat) and data from these electrodes was excluded from the analysis. Figure is adapted from Fig. 5A in McGarrity et al. (2017).

All data were processed in MATLAB (R2016a) (The MathWorks, Inc., USA) using the FieldTrip toolbox (Oostenveld, Fries, Maris, & Schoffelen, 2011). To reduce noise, far-field effects and shared signal from all the electrode recordings, a montage of seven bipolar channels was digitally created using a pairwise bipolar subtraction (Land et al., 2012) of the eight original electrodes. Data was then pre-processed in fieldtrip by re-referencing with a common reference over all bipolar channels (except for the functional connectivity analysis) to remove any remaining linear gradient affecting all bipolar pairs, DC offset removal for each episode, and a band-stop filter between 49.5 and 50.5 Hz to remove the electric line-noise. The most medial and/or the most lateral electrodes of the 2-mm 8-electrode recording array were located outside of the hippocampus (typically 1–3 electrodes per rat) (**Figure 2**, bottom) and our previous multi-unit analysis revealed that hippocampal picrotoxin infusion did not affect any of the multi-unit parameters analysed from these electrodes (consistent with the densely packed fibre bundles surrounding the hippocampus) (McGarrity et al., 2017). If both of the electrodes used to calculate a synthetic bipolar electrode were outside of the hippocampus, that particular bipolar channel was removed, otherwise it was included in the analysis. As bipolar subtraction was not appropriate for multi-unit activity, in order to match the 7 bipolar channels used for the LFP analysis, electrodes 2 to 8 were selected and matched to their corresponding bipolar channel (i.e., electrode 2 to bipolar 1, electrode 3 to bipolar 2 etc.). Electrodes 2 to 8 were selected in place of 1 to 7 because in most cases electrode 1 (the most medial) sat outside the hippocampus and had to be excluded anyway.

### 2.2 Semi-automated separation of LFP states

The hippocampal LFP recordings under isoflurane anaesthesia clearly showed the three distinct LFP states outlined in the Introduction, including a burst, suppression and continuous state (**Figure 1A**). The simultaneously recorded multi-unit activity also appeared to vary depending on the LFP state, with the burst state tending to show a higher multi-unit firing than the suppression state, and the continuous state characterized by continuous firing at a similar rate to that in the burst state. Our semi-automated separation strategy was to start by separating the combined burst-suppression state from the continuous state and then to separate the burst from the suppression state.

Simple thresholding based on continuous amplitude/power measures did, with our data, not result reliably in state separation that corresponded to the states that would be assigned by visual inspection (although thresholding worked for the example LFP traces in **Figure 1A**, it did not reliably separate the LFP states when these showed less pronounced differences; for examples, see **Figure 1B**, right). Therefore, we used a higher order moment (kurtosis) of the distribution of peak amplitude values (rather than the power of the continuous signal). To separate the three states, for each rat the channel with the most marked difference between LFP states based on visual inspection of the raw LFP traces, the ‘LFP state-defining’ channel, was selected (see **Figure 1B** for example). For this LFP state-defining channel, the peaks and troughs of the LFP trace were identified over baseline and post-infusion periods, by identifying the sign change of the first derivative. The distribution of these peaks and troughs identified from the sign-change turned out to be unimodal (not shown), and thus not enabling a clear separation into discrete states. In contrast, a measure of the kurtosis (‘tailedness’) of the amplitude distribution of those peaks and troughs separated the burst-suppression and continuous states (**Figure 3**). The kurtosis was calculated for a 10-s sliding window with a 1-s step size (**Figure 4A,B**). A histogram of kurtosis values (a total of 4391) was plotted in 100 bins (**Figure 4C**), and the most distinct local minimum (the first or second) separating peaks in the multimodal distribution, was selected as threshold to separate the combined burst-suppression states from the continuous state (**Figure 4D**). If kurtosis values exceeded this threshold, the corresponding time points were marked as being in the ‘burst-suppression’ state, whereas time points with subthreshold values were marked as in the ‘continuous’ state (**Figure 4E**). These labels were then applied to the samples of every channel from the same rat.

**Figure 3:**
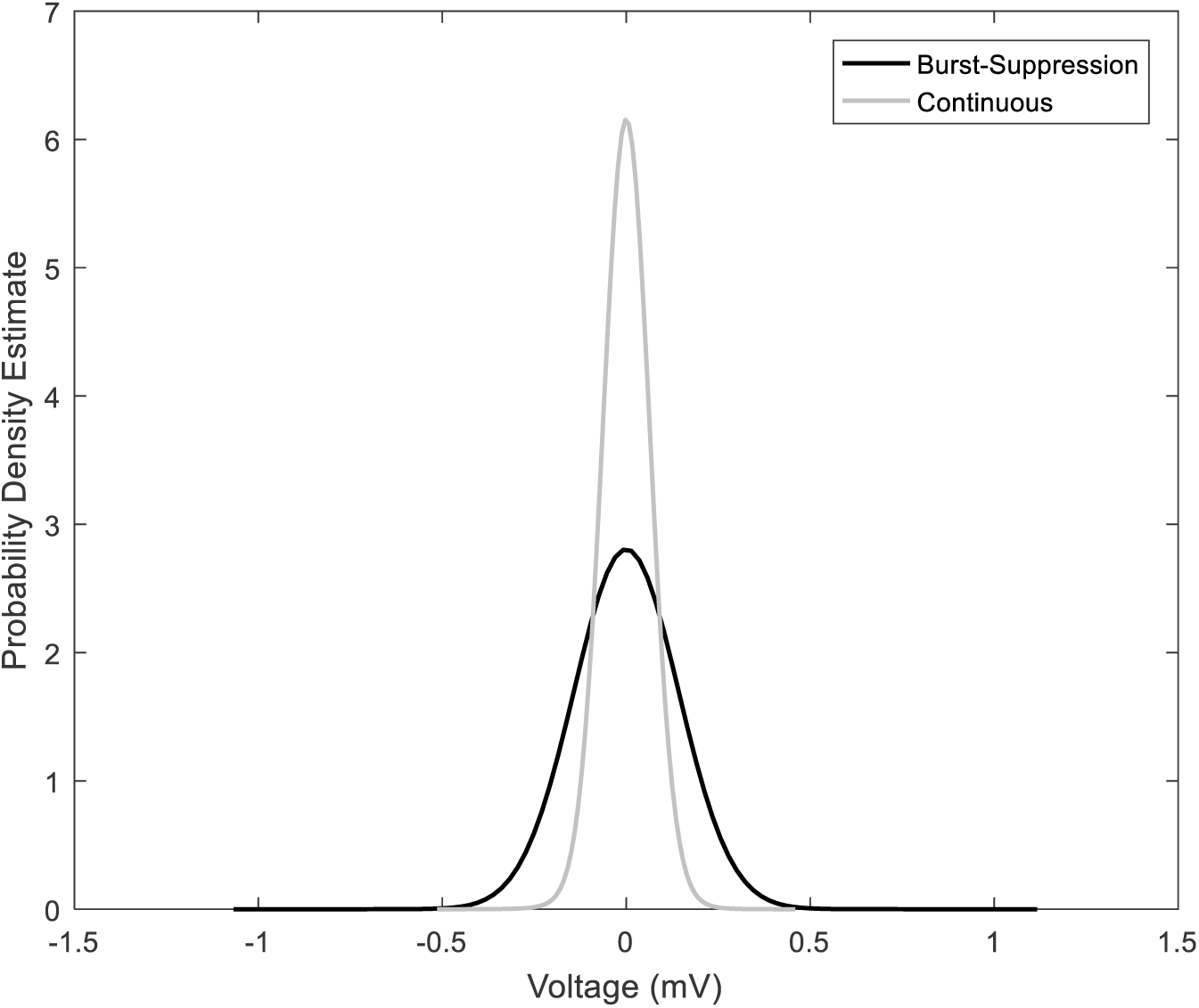
The distribution of amplitude of peaks and troughs in the burst-suppression state and the continuous state differ in kurtosis, but not modality. Using a kernel distribution, a probability density estimate was calculated for the amplitudes of peaks and troughs based on visually identified 100-s segments belonging to the combined burst-suppression or the continuous state, from the 400-s trace shown in Figure 1. Both states show a unimodal amplitude distribution, but the burst-suppression state shows a higher kurtosis (tailedness).

**Figure 4:**
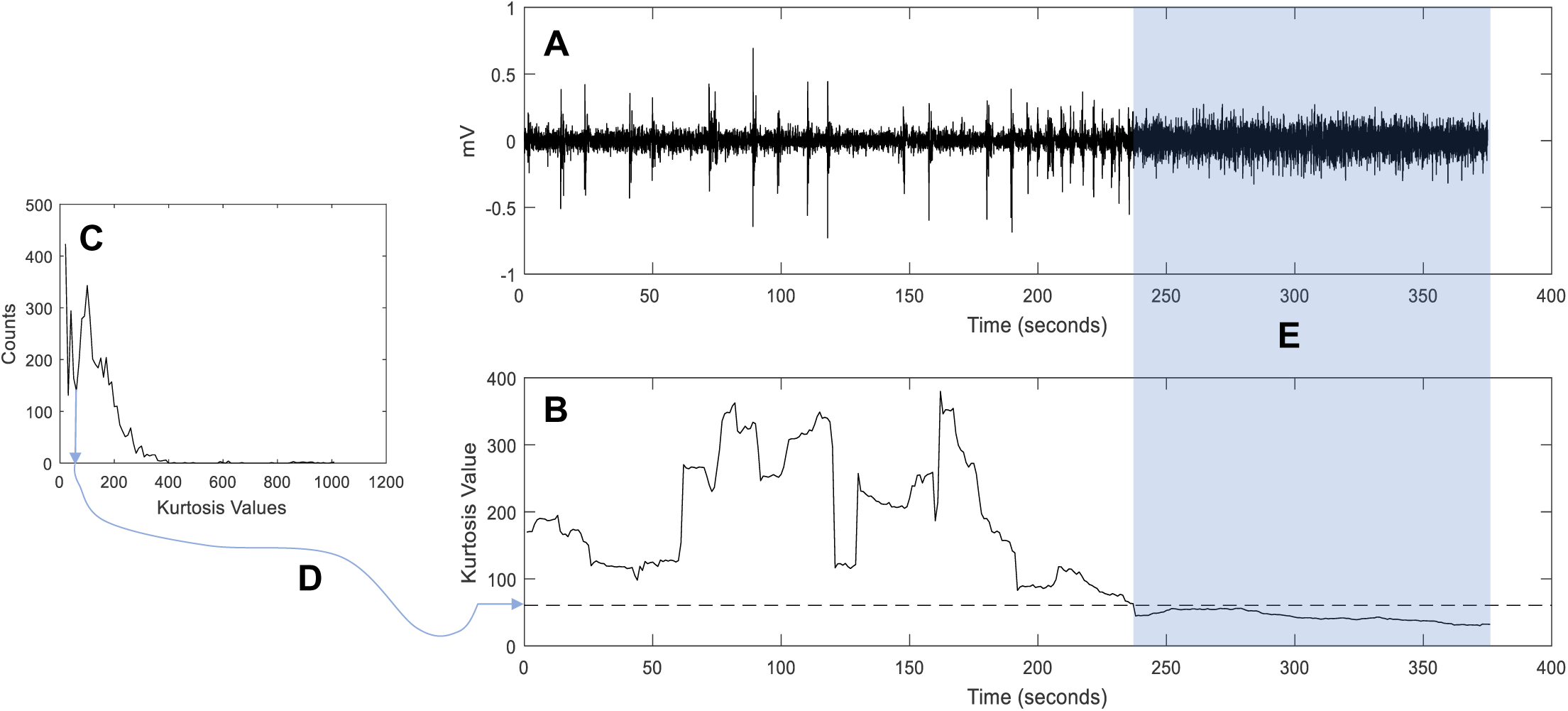
Semiautomatic separation of burst-suppression and continuous LFP states, using the kurtosis values of the distribution of the peak and trough amplitudes of the LFP trace. **A** 400-s LFP trace, clearly showing separate burst-suppression (white background) and continuous states (shaded) (same 400-s trace as shown in Figure 1). **B** The time course of the kurtosis values of the 400-s LFP trace, calculated using a 10-s long, 1-s sliding window. **C** Histogram of the distribution of kurtosis values over baseline and post-infusion data (i.e., the whole recording period, not only the 400-s period presented in **A** and **B**). **D** In this case, the second minimum in the histogram (**C**) corresponds to the kurtosis threshold value (stippled line in **B**) that separates the burst-suppression and the continuous state.

In a final step, we further separated the ‘burst-suppression’ state into the burst and suppression components, using a semiautomatic LFP-amplitude thresholding routine in Fieldtrip. To this end, a distribution of the amplitude envelope of the analytic signal was formed. Samples that exceeded a z-value of 5 and those falling within a symmetrical 0.25s window around the threshold crossing were consecutively labelled as ‘Burst’ state. This thresholding was run on each individual channel to prevent summation of z-values, which could occur from synchrony across channels and convolute the routine. The output of this routine gave the boundaries between low amplitude activity, corresponding to the suppression state, and the burst state.

Not all rats were spending time in each of the LFP states for every 5-min bin of the recording period, and four rats were not spending any time in the continuous state during baseline (therefore not allowing baseline normalisation of the post-infusion data). Therefore, the number of rats contributing to the different analyses of state-separated data could differ from the overall number of rats from which recordings were made (**Figure 5**). The reduced sample size during the continuous state implies that the statistical power of any analysis in the continuous state is reduced compared to the analysis of the other two states.

**Figure 5:**
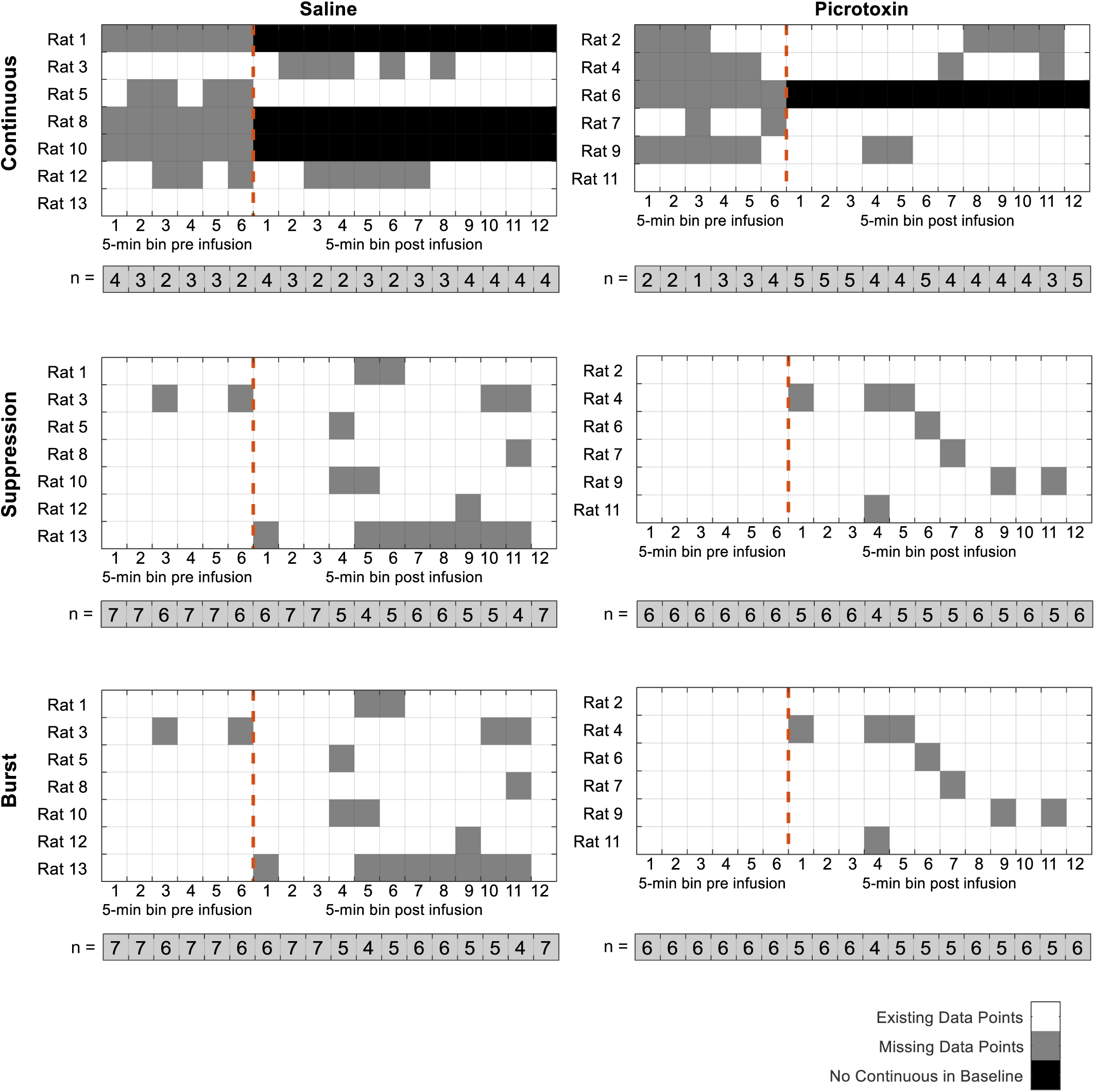
State-separated data available in saline and picrotoxin infusion groups for each of the 6 baseline and 12 post-infusion 5-min bins. For each of the drug (saline or picrotoxin)-state combinations a grid is shown to indicate missing data points, with the time bin on the x-axis and the rat ID on the y-axis. Underneath is the total number n of rats that contributed values for each 5-min bin in the different drug-state combination.

### 2.3 State-separated analyses of hippocampal LFP properties

#### 2.3.1. Power-spectral density

Power at different frequencies was analysed separately for the different LFP states in FieldTrip, using Fast Fourier Transform with a Hanning window from 0.5 to 40 Hz. These power-spectra were subsequently averaged across channels and compared across the different states. Given the substantial raw-power differences between the different states (already separated in this analysis) and in order to investigate potential differences in relative spectral density, for each individual channel we additionally normalized the spectra to the total area under the curve of the power spectrum before comparison across states.

#### 2.3.2. Connectivity

Phase-Lock Values (PLV) were calculated to indicate the phase-coherence (a measure of neuronal synchronization that is independent of amplitude correlations) of two signals originating from a pair of electrodes (measuring local activity due to their bipolar montage) (Aydore, Pantazis, & Leahy, 2013; Bastos & Schoffelen, 2015; Srinivasan, Winter, Ding, & Nunez, 2007). PLVs were calculated for every possible pair of hippocampal electrodes for every 0.5-Hz frequency bin. Because the individual electrodes between rats were not at consistent homological locations within the temporal to intermediate hippocampus, instead of averaging PLVs of electrode-pairs across subjects (as is common in EEG/MEG analysis where a standard mapping of electrodes does exist e.g. (Oostenveld & Praamstra, 2001), we used a different method to extract the communality of connectivity between all electrode pairs and to then calculate a statistic of this across rats (Cohen, 2015). To this end, we used singular value decomposition (SVD) over the imaginary part of the PLV matrix of all states and times. We used the imaginary part of the complex phase-locking-value (complex cross-spectrum normalized on power in individual trials) as this is not corrupted by volume-conduction problems (Nolte et al., 2004). For each rat and frequency bin, we extracted the singular vectors U and V that satisfy the equation PLV = U * S * V’, with S being the matrix of singular values and PLV the matrix of imaginary PLVs. The individual imaginary PLV matrices for each state and time combination were then projected into this space using the respective singular vectors and the first two “singular values” were extracted. These two values represent the connectivity strength within the two dominant ‘neural networks’ of the rat in the area of the temporal to intermediate hippocampus sampled by the bipolar 7-electrode montage.

#### 2.3.3. Multi-unit activity (MUA)

We compared the following key MUA parameters between LFP states: overall firing rat, bursts per minute, percentage of spikes in burst and burst duration. The original MUA analysis in our previous study (McGarrity et al., 2017) had been completed using NeuroExplorer (version 4). However, in order to examine the state-dependency of MUA parameters (and, subsequently, also of the previously reported disinhibition-induced enhancement of MUA burst parameters, see below), we replicated the original analysis using MATLAB, where we could then separate the different snippets of MUA by their associated LFP state. The MUA data was separated according to the different LFP states based on their time stamps in a way that preserved the original MUA bursts, with each burst assigned to the state occupied longest during the burst duration.

#### 2.3.4. Statistical comparison of the three distinct hippocampal LFP states during baseline

To analyse the effect of the different states on the overall LFP and MUA we compared the baseline data, removing the factor of drug. For the baseline frequency specific effects (power-spectra and PLV eigenvalues) we compared each state to both other states in a separate analyses, after running a moving average over the frequency values, with a kernel of 1.5Hz sliding over 0.5Hz. The statistical analysis of these effects were carried out using pairwise comparisons at each frequency bin. In order to address the multiple comparisons problem arising from the simultaneous analysis of 80 different frequency bins, we conducted a cluster permutation analysis (Hayasaka, Phan, Liberzon, Worsley, & Nichols, 2004; Maris & Oostenveld, 2007). This method runs a mass-univariate independent samples t-test for each frequency bin, for both real data pairs (of the conditions to be compared) and randomly assigned pairs (combining data of both conditions to obtain a reference distribution). Adjacent frequency bins that pass the univariate significance threshold form a cluster and a summary statistic of these clusters (e.g. summed t-value) is calculated for real data and reference distribution data. Only those clusters from the comparisons of the real data pairs which cut-off less than 5% of the maximum cluster-statistic from the reference distribution (random pairs) were considered as statistically significantly different.

For the state-separated analysis of MUA, a problem of missing data points over time arose due to the limited time each rat spent in each state (see **Figure 5** for the missing data in different 5-min bins). Therefore, it was not possible to conduct a repeated-measures ANOVA and we used a pooling strategy. More specifically, we used the 5-min bins in the different states as the units of observation, such that data from all rats and time bins were pooled according to state – after averaging across channels (for each rat). This was then subjected to a between-subjects one-way ANOVA, with LFP state as the between-subjects factor; this reflects a fixed-effect analysis.

### 2.4 Analyses of LFP state-dependent drug infusion effects

#### 2.4.1 Power-spectral Density

For the comparison of frequency-specific effects of drug infusion, the difference between pre- and post-infusion power was expressed as log of the ratio between post-infusion (power-spectra over 60 min) and pre-infusion (baseline; power spectra over 30 min) values for each channel. These difference values were then averaged per rat for statistical analysis, then for each drug group for plotting. The statistical analysis of the frequency specific effects of picrotoxin compared to saline infusions on LFP power were carried out using cluster permutation where, separately for each LFP-state, the log-ratio of both drug groups was compared.

#### 2.4.2 Connectivity

The first two PLV ‘Eigenvalues’ for each rat (see State-separated analysis of hippocampal LFP properties, Connectivity) were averaged to give a single measure of aggregate connectivity. To normalise post-infusion values to pre-infusion (baseline) values, the pre-infusion values were subtracted from the post-infusion values. As was done for the spectral power analysis, the cluster permutation statistic was used to compare the effects of picrotoxin vs. saline for each of the LFP-states separately.

#### 2.4.3 MUA

Our previous analysis, which did not separate MUA data according to LFP state, showed that hippocampal disinhibition by picrotoxin causes enhanced burst firing, as reflected by increased bursts per minute, percentage of spikes in burst, and burst duration (McGarrity et al., 2017). Therefore, our present analysis focused on these burst parameters.

To assess the effect of picrotoxin on the state-separated MUA analysis, we could not apply the mixed-model ANOVA of MUA parameters with drug as between-subjects and time (5-min bin) as repeated-measures factors, which we used in our original MUA data analysis (which did not separate data by LFP state) (McGarrity et al., 2017). This is because of the many missing data points, following separation of data according to LFP state (see **Figure 5**). Therefore, we used the same pooled analysis as for the comparison of MUA parameters between LFP states during baseline, however, here with the additional factor of infusion group. Again, data were pooled over time and rats and separated according to LFP-state and infusion group. Prior to pooling, data for each rat, in each state, were normalised by subtracting the average of the pre-infusion values for each channel from the individual post-infusion values for the corresponding channel; subsequent to this, an average was taken across the channels, for each rat and time point separately. The normalised and pooled MUA data were subsequently analysed using a two-way ANOVA, with LFP state and infusion condition as between-subjects factors (and time-bins and rats as units of observation). A significant infusion condition X state interaction would statistically support state-dependent drug effects.

## 3. RESULTS

### 3.1. Three distinct hippocampal LFP states under isoflurane

#### 3.1.1. Differences in frequency-dependent power and functional connectivity

To compare quantitatively the three distinct hippocampal LFP states (**Figure 1A**), we combined the baseline data (i.e., data recorded before any drug infusion) from all infusion groups. During the baseline period, all rats showed the burst and suppression states, but not all rats showed a continuous state, resulting in a sample size of n=13 rats for the burst and suppression state and of n=9 rats for the continuous state (**Figure 5**).

The three LFP states substantially differed with respect to overall raw power and with respect to relative power at different frequencies (**Figure 6**). The overall raw power in the burst state was significantly and substantially higher than in the suppression (cluster level p = 0.000999, corrected for multiple comparisons; cluster spanning across all frequencies, i.e. 1-39 Hz) and continuous states (p = 0.005, cluster including 1-22.5Hz). Suppression and continuous states did not differ significantly (p>0.05) indicating similar overall power, although at lower frequencies power tended to be somewhat higher in the continuous compared to the suppression state (**Figure 6A**). With respect to relative power at different frequencies, the burst state showed a higher proportion of power within delta/low theta than the other two states and tended to show a lower proportion of power in high frequencies from about 20 to 40 Hz than both the suppression and continuous states, which were both characterized by conspicuous gamma ‘bumps’ in the normalised power spectra in this frequency range (**Figure 6B**). For the comparison between burst and continuous states, a significant cluster was found at 3.5-10 Hz (p = 0.024) and between burst and suppression states a significant cluster was found at 4-14 Hz (p = 0.009). The numerical differences at higher frequencies did not reach statistical significance; neither for the comparisons of continuous and burst states (p = 0.16) nor suppression and burst states (p = 0.10).

**Figure 6:**
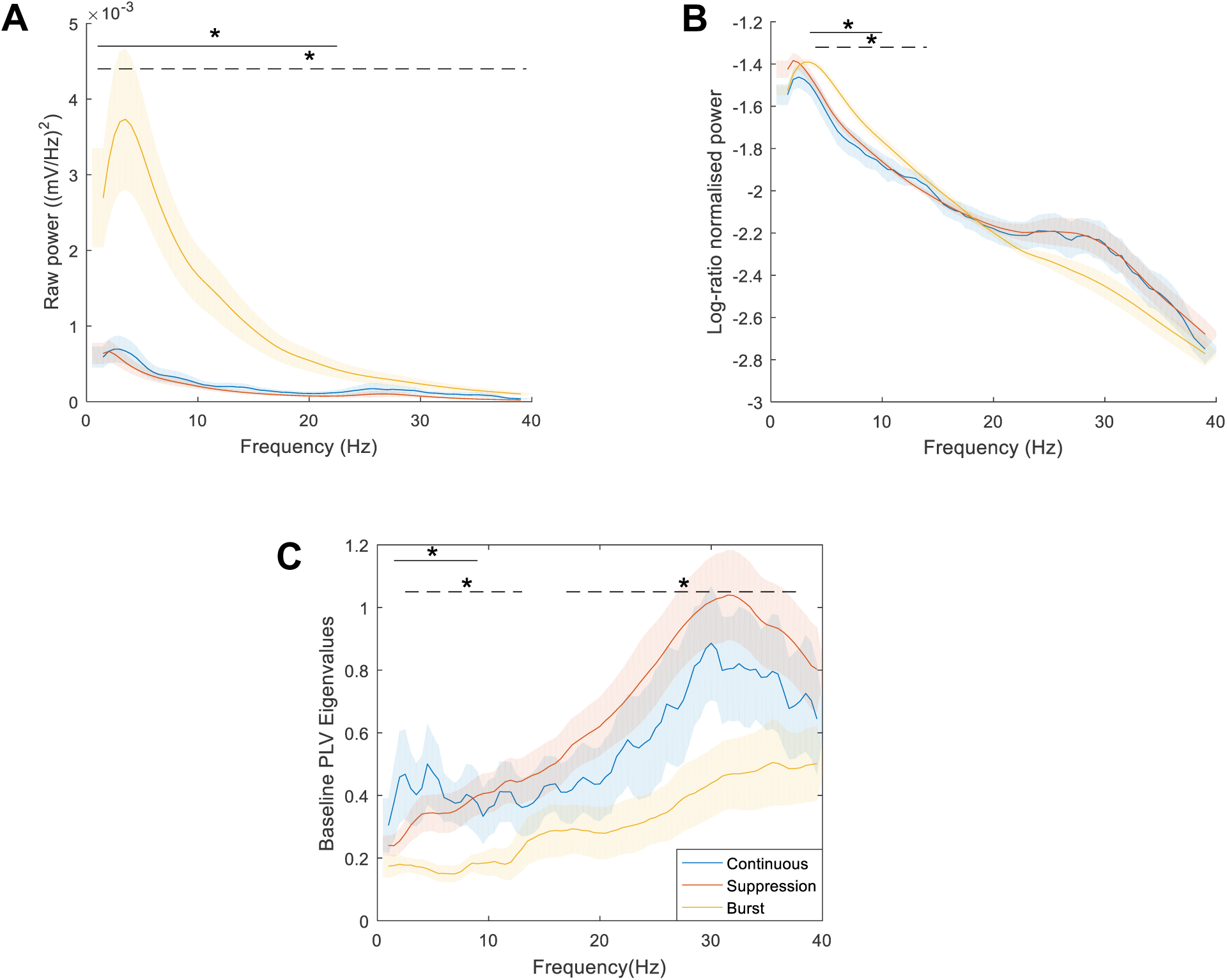
The three hippocampal LFP states – differences in frequency-dependent power and connectivity. (**A**) Raw power spectral density, (**B**) log-ratio normalised power (representing relative power at different frequencies), and (**C**) phase-lock eigenvalues are shown across frequencies for each of the three LFP states. All values shown as mean±SEM. The frequency ranges showing significant (p<0.05) differences based on cluster permutation statistics are indicated by an asterisk (*) and a solid line for comparisons between burst and continuous, and a dashed line for comparisons between burst and suppression states. Number of rats contributing data to the different states: burst, n = 13; suppression, n = 13; continuous, n = 9.

Regarding connectivity, we computed an aggregate measure of phase-locking between all electrode pairs spread across the temporal to intermediate hippocampus (see Methods). To exclude any contamination by volume conduction, only the imaginary part of the PLVs was used and submitted to a SVD, of which we report the first two diagonal (Eigen-)values. Across the frequency range examined (0.7 to 40 Hz), the burst state showed generally lower connectivity than both suppression and continuous states, which showed very similar connectivity (**Figure 6C**). One significant cluster was found for the comparison between burst and continuous states at 1.5-9 Hz (p=0.018), and two significant clusters for the comparison between burst and suppression states: one cluster at 2.5-13 Hz (p =0.019) and another cluster at 17-38 Hz (p = 0.005). In all these clusters connectivity was reduced in the burst state, in comparison to both the continuous and the suppression state.

Overall, in terms of total power, relative power at different frequencies and functional connectivity, the burst state of the hippocampal LFP markedly differs from the suppression and continuous states, whereas the latter two states show similar characteristics in these parameters.

#### 3.1.2. Differences in associated MUA

At baseline, some of the MUA parameters differed between states (**Table 1**). The pooled analysis explained in the methods revealed that states significantly differed with respect to overall firing rate (spikes per second) (F(2,180) = 3.37, p = 0.037), with the firing rate in the burst state being higher or tending to be higher, respectively, than in the suppression state (p = 0.02) and continuous state (p = 0.058), which did not differ (p = 0.904). States also differed significantly with respect to percentage spikes in burst (F(2,177) = 9.44, p < 0.0001), with both burst and continuous states, which did not differ significantly (p = 0.108), showing a higher percentage of spikes in bursts than the suppression state (p = 0.002 and p<0.0001, respectively). States did not differ significantly with respect to bursts per minute (*F*(2,177) = 0.95, p = 0.383) or burst duration (*F*(2,172) = 1.19, p = 0.307). Overall, this quantitative comparison of MUA parameters is consistent with the impression based on visual inspection of the MUA data in the three states (**Figure 1A**) that MUA activity is ‘suppressed’ in the suppression state compared to the burst and continuous state.

**Table 1:** Baseline values for different MUA bursting parameters in the three LFP-states. Mean ± SEM. * indicates significant difference between states (p < 0.05), with the number indicating the pair of states that differ.

|  | **Bursts per minute** | **% spikes in burst** | **Mean Burst duration (s)** | **Spikes per second** |
| --- | --- | --- | --- | --- |
| **Continuous** | 441.50 ± 65.95 | 35.10 ± 2.22*<sup>2</sup> | 0.0094 ± 0.00025 | 21.29 ± 5.12 |
| **Suppression** | 325.85 ± 47.75 | 20.26 ± 1.96 *<sup>1</sup> *<sup>2</sup> | 0.0088 ± 0.00039 | 24.06 ± 3.16*<sup>1</sup> |
| **Burst** | 380.32 ± 44.03 | 29.26 ± 2.14*<sup>1</sup> | 0.0093 ± 0.00041 | 65.33 ± 18.86*<sup>1</sup> |

### 3.2. Hippocampal neural disinhibition did not affect the expression of the three hippocampal LFP states

During the post-infusion period, the cumulative time spent in the different LFP states was similar in rats infused with picrotoxin and saline (**Figure 7**). Independent sample t-tests revealed that there was no significant difference between the picrotoxin and saline groups in the total time spent in any of the three LFP states (continuous state, t(7)<1; suppression and burst state: t(11)<1).

**Figure 7:**
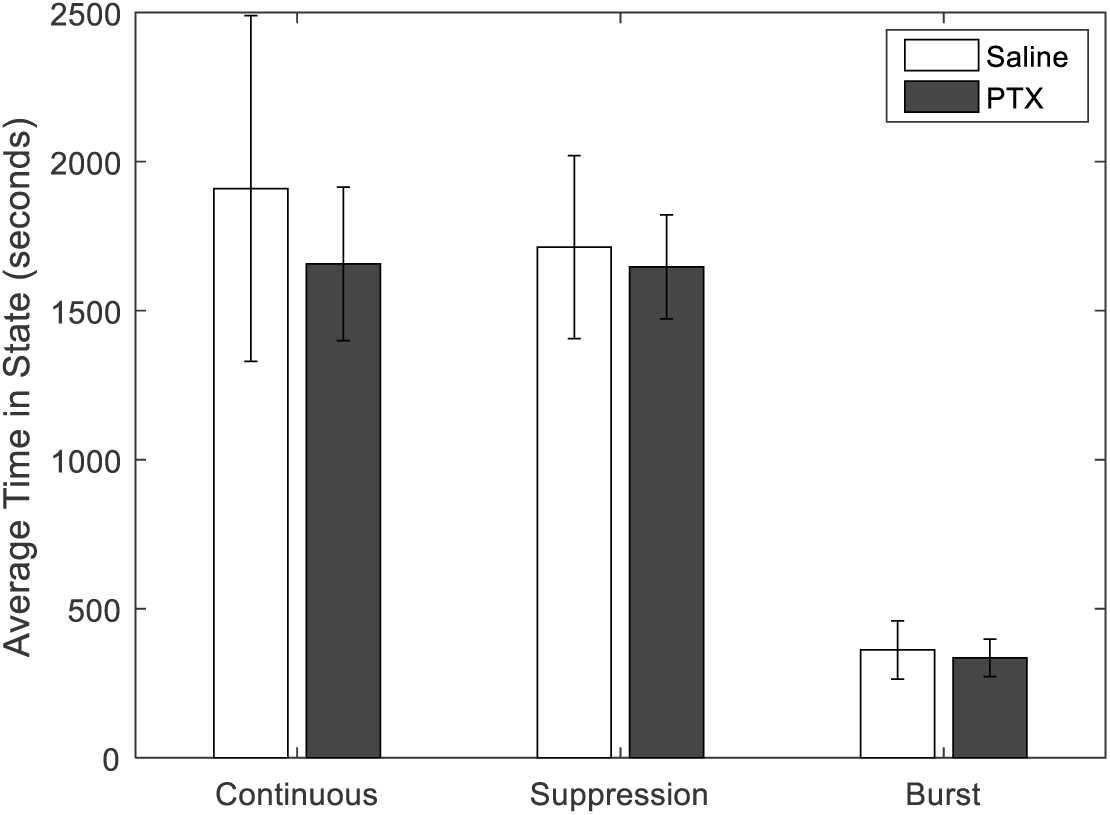
Hippocampal neural disinhibition does not affect the expression of the three hippocampal LFP states. Cumulative time (mean ± SEM) spent in each LFP state by the two drug groups during the post-infusion period.

### 3.3. Hippocampal neural disinhibition caused state- and frequency-dependent effects on hippocampal LFP power and functional connectivity

Hippocampal picrotoxin infusions, compared to saline, tended to increase power in the lower frequencies (<20 Hz) and to decrease power in the higher frequencies (>20 Hz), in both burst and suppression states, while having minimal effect in the continuous state (**Figure 8A,C,E**). However, only the picrotoxin-induced increase in power at 6-18Hz in the burst state attained statistical significance (p = 0.017). In the continuous state, power was numerically higher during the post-infusion period, as compared to the pre-infusion baseline; this was regardless of infusion, as indicated by a positive difference in log ratio power, especially at lower frequencies, which indicates ‘baseline drift’. There was no difference between the drug groups in this state (no clusters) (**Figure 8A**).

Hippocampal picrotoxin tended to decrease functional connectivity (as reflected by phase-lock Eigenvalues) at higher frequencies in the burst and suppression states and at lower frequencies in the continuous state (**Figure 8B,D,F**). However, the picrotoxin-induced decreases in connectivity reached statistical significance only in the burst state, in a high-frequency cluster ranging from 29.5-33 Hz (p = 0.049), but there was no significant picrotoxin-induced decrease in connectivity during the suppression state (no clusters). The visually apparent decrease in the continuous state (at lower frequencies) also failed to reach statistical significance, although there was a trend in a cluster from 6-7 Hz (p = 0.065).

**Figure 8:**
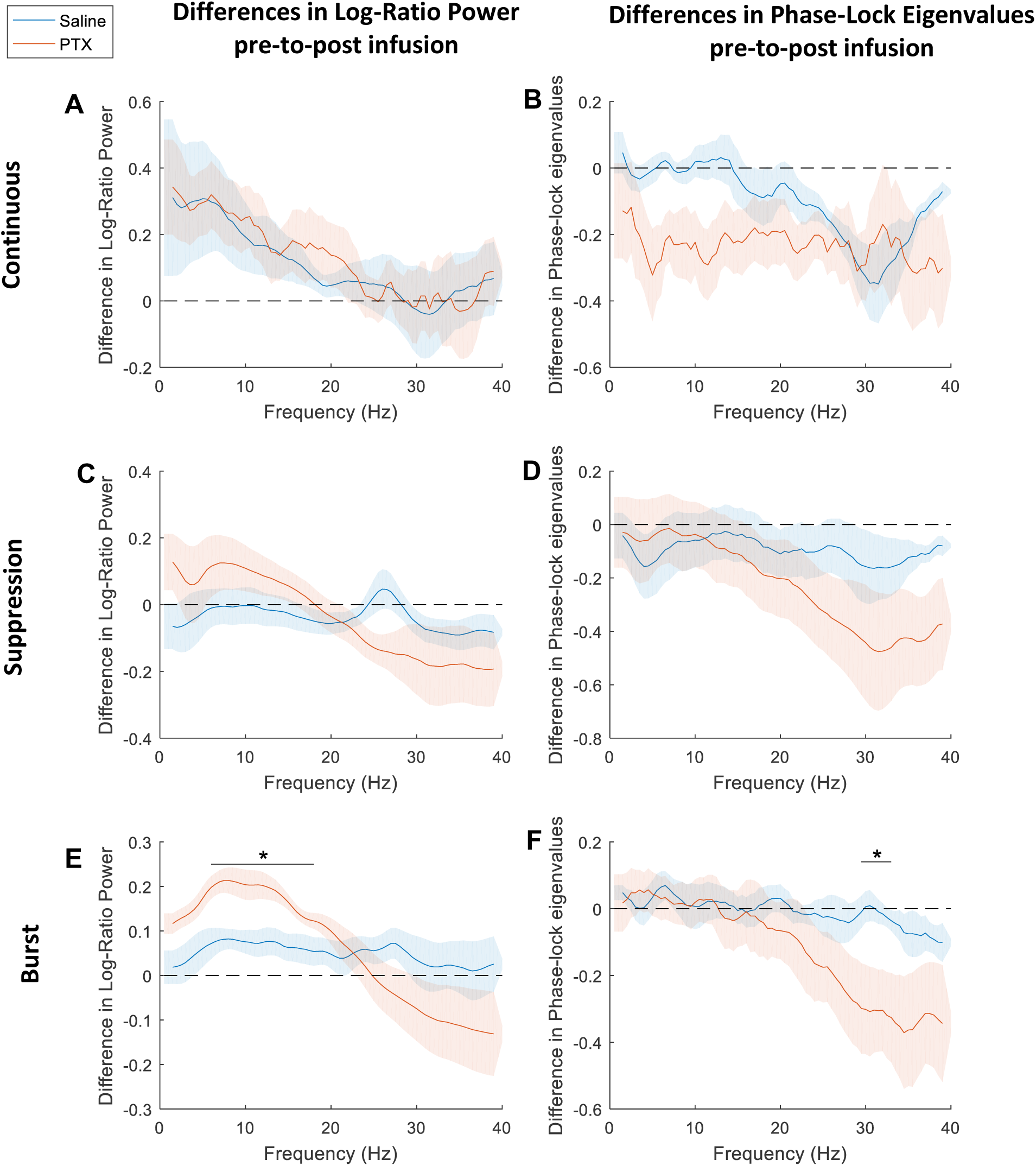
Hippocampal neural disinhibition causes state- and frequency-dependent effects on hippocampal LFP power and functional connectivity. Frequency-dependent impact of hippocampal picrotoxin or saline infusion on power (left) and on functional connectivity (right) in the continuous (top panel), suppression (middle panel) and burst (bottom) states. The difference in power between post-infusion and pre-infusion (baseline) periods after saline and picrotoxin infusions (left) was compared by plotting the log of the ratio between post-infusion and pre-infusion power for both infusion groups across frequencies and separately for each state (**A, C, E**). The difference in functional connectivity between post-infusion and pre-infusion (baseline) periods after saline and picrotoxin infusions (right) was compared by plotting the difference between post-infusion and pre-infusion phase-lock Eigenvalues for both infusion groups across frequencies and separately for each state **(B, D, F).** All values shown as mean±SEM. The dotted line (value of 0) represents no change from the pre- to the post-infusion period, while positive numbers indicate an increase from baseline and negative numbers a decrease from baseline in the post-infusion period. The frequency ranges showing significant (p<0.05) differences between the picrotoxin and saline infusion groups based on cluster permutation statistics are indicated by an asterisk (*) and a solid line.

### 3.4 State-dependence of enhanced multi-unit burst firing caused by hippocampal neural disinhibition

We first replicated the picrotoxin-induced increases in burst parameters, including burst per min, percentage of spikes in bursts and burst duration, as calculated by the Neuroexplorer software and originally reported by McGarrity et al. (2017), using a MATLAB code that could be run with the segmented LFP state-separated data (**Figure 9A,B,C**); the original neuroexplorer data for rats that were excluded from the LFP analysis (see Methods 2.1.) was inputted directly into MATLAB to complete the dataset for comparison of analyses (i.e., n=8 rats in the picrotoxin group and n=7 rats in the saline group). Next we plotted the time course of the multi-unit burst parameter following picrotoxin and saline infusions separately for the three different LFP states, using the state-separated multi-unit data (**Figure 9 D-L**) (with sample sizes for the different 5-min bins indicated in **Figure 5**). Visual inspection of these plots indicates that, numerically, the picrotoxin-induced increase of bursts per min appears to be weaker during the continuous LFP state (**Figure 9D-F**) compared to the burst state (**Figure 9G-I**) and suppression state (**Figure 9J-L**). In order to examine directly the LFP state-dependence of the picrotoxin effects, we pooled the time-deconstructed post-infusion values of the three multi-unit burst parameters, subtraction-normalised to baseline, and separated them by infusion group and LFP state (**Figure 9M-O**). ANOVA of these values, using LFP state and drug infusion as independent variable (see Methods 2.4.3.), revealed a significant main effect of drug infusion for all three burst parameters (outcomes of statistical analysis not shown), reflecting that, overall, picrotoxin infusion increased these values as compared saline infusion. Importantly, the ANOVA of burst duration (**Figure 9O**) revealed a significant interaction of drug infusion X LFP state (F(2,316)=3.86, p=0.022), reflecting that picrotoxin increased burst duration in the burst and suppression states, but not the continuous state. Although, numerically, the picrotoxin-induced increases in bursts per min and percentage of spikes in bursts also tended to be somewhat weaker in the continuous, compared to the other two states (**Figure 9M,N**), ANOVA did not support significant interactions of drug infusion X LFP state for these two parameters (burst per min: F(2,343) = 1.118, p = 0.328; percentage spikes in bursts: F(2,343) <1). Overall, the increased multi-unit burst duration following hippocampal neural disinhibition was markedly less pronounced in the continuous state of the hippocampal LFP, as compared to the other two states. The other two burst parameters showed numerical tendencies in the same direction, but these were not statistically significant.

**Figure 9:**
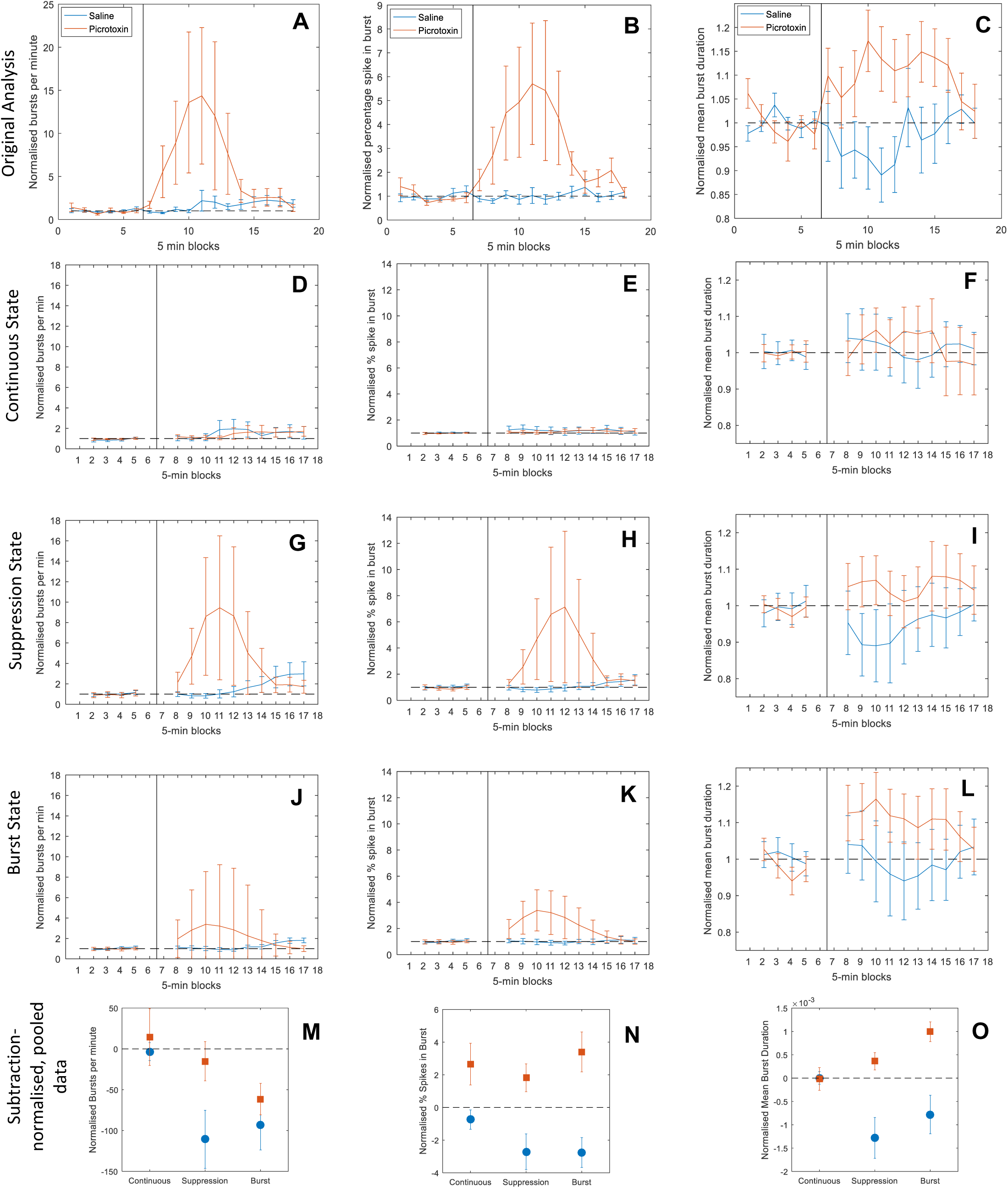
State-dependence of enhanced multi-unit burst firing caused by hippocampal neural disinhibition. The top row displays a replication of the time course of multi-unit burst parameters during baseline and following infusion of picrotoxin and saline, across all LFP states, i.e. without state separation, as originally reported by McGarrity et al. (2017), but carried out in Matlab rather than Neuroexplorer v4. Hippocampal neural disinhibition by picrotoxin markedly enhanced hippocampal multi-unit burst firing as compared to saline infusions, as reflected by increases in (**A**) bursts per minute, (**B**) percentage spikes in burst, and (**C**) burst duration (and statistically supported by a significant infusion X time interaction, McGarrity et al., 2017). All values are division-normalised to the baseline value and mean±SEM. The three rows underneath display the same time course analysis, but separated by LFP state, using the Matlab sripts developed in the present study: (**D,E,F**) continuous state; (**G,H,I**) suppression state; and (**J,K,L**) burst state. Visual inspection indicates that the picrotoxin-induced enhancement of burst parameters is less pronounced in the continuous, compared to the other two states. To visualise the potential interaction of hippocacampal infusion and LFP state, the bottom row displays the time-deconstructed and pooled post-infusion values (subtraction-normalised to baseline, mean ±SEM) of (**M**) burst per min, (**N**) percentage spikes in bursts, and (**O**) burst duration, for the picrotoxin and saline infusion groups and separated by LFP state; these plots also indicate that the picrotoxin-induced increase in burst parameters is, at least numerically, less pronounced in the continuous than in the other two states.

## 4. Discussion

We, first, developed a semi-automated method based on the kurtosis of the LFP peak-amplitude distribution and amplitude envelope thresholding to separate three distinct hippocampal LFP states in the isoflurane anaesthetized rat: burst, suppression and continuous states (**Figure 1**), and we described some of the properties of these three LFP states. The burst state shows substantially larger overall raw power across the frequency range examined, compared to the suppression and continuous states (**Figure 6A**), as well as higher relative power in the delta/theta range (<15 Hz), but lower relative power in the high beta/low gamma range (>20 Hz) (**Figure 6B**). Moreover, compared to suppression and continuous states, the burst state showed lower frequency specific ‘functional connectivity’ across the recording area in the temporal to intermediate hippocampus, as reflected by reduced phase-lock eigenvalues of the LFP signals recorded from the different electrodes (**Figure 6C**). In terms of MUA, overall neuronal firing was higher in the burst state than in the two other states, whereas the percentage of spikes fired in bursts was higher in both the burst and continuous states as compared to the suppression state (**Table 1**). Finally, the expression of the hippocampal LFP states, as reflected by the cumulative duration of these states across the recording period, was not affected by hippocampal neural disinhibition (**Figure 7**).

Second, and importantly, we then showed that hippocampal neural disinhibition by picrotoxin changed the hippocampal LFP in a state- and frequency-dependent way (**Figure 8**). In both burst and suppression states, but not the continuous state, hippocampal neural disinhibition tended to increase LFP power at lower frequencies (<20 Hz) and to decrease power and functional connectivity at higher frequencies (>20 or >15 Hz, respectively). There was also a trend towards picrotoxin decreasing functional connectivity in the theta range (6-7 Hz) during the continuous state.

Third, we also found that the increase in multi-unit burst parameters by hippocampal neural disinhibition, which we reported in a previous study (McGarrity et al., 2017), was partially dependent on the hippocampal LFP state: disinhibition-induced increases in burst duration were limited to the burst and suppression state and not apparent in the continuous state (**Figure 9**).

### 4.1. Three distinct hippocampal LFP states under isoflurane anaesthesia

Visual inspection of the LFP recorded from the temporal to intermediate hippocampus in isoflurane-anaesthetised rats indicates three distinct LFP states, including burst, suppression and continuous states (**Figure 1A**). We found that the kurtosis of the LFP peak and trough amplitude distribution was the best metric to separate the periods including burst and suppression states from the continuous state. Using this kurtosis measure, we thus separated the continuous state from the combined burst and suppression states, which could then be separated by amplitude envelope thresholding. Power analysis corroborated the impression based on visual inspection of the LFP traces, that raw power during the burst state is substantially higher than in the other two states, owing to large amplitude LFP bursts, and that the continuous state tends to show higher power than the suppression state (although the latter difference failed to reach statistical significance). Interestingly, analysis of relative power revealed a marked beta/gamma shoulder (24 to 34 Hz) in the power spectra of continuous and suppression states which was absent in the burst state. Others have previously reported similar beta/gamma shoulders in the hippocampal LFP during continuous-like LFP states under anaesthesia (Land et al., 2012; Pagliardini, Gosgnach, & Dickson, 2013). However, if the burst state is not separated from the other LFP states, hippocampal LFP power spectra recorded under anaesthesia do typically not show a beta/gamma shoulder (Clement et al., 2008; Lustig et al., 2016; Wolansky et al., 2006; and our unpublished observations). This likely reflects that the power spectra are dominated by the large-amplitude burst state (which lacks a beta/gamma shoulder). Overall, in the frequency range examined, continuous and suppression states tended to show higher connectivity, as indicated by phase-lock Eigenvalues, than the burst state across the recording area in the temporal to intermediate hippocampus. In the continuous state, a peak in connectivity was apparent at lower frequencies (<10 Hz, **Figure 4C**), which included the theta range, although statistically connectivity in this range did not significantly differ between continuous and suppression states. The numerically increased theta-range connectivity may correspond to findings of characteristic activity in the theta range in related hippocampal LFP states identified by others. More specifically, several studies have described hippocampal LFP states under anaesthesia, distinct from periods characterized by burst-suppression patterns and referred to as ‘continuous’/ ‘activated’ (Clement et al., 2008; Wolansky et al., 2006) or ‘light anaesthesia’ (Land et al., 2012; Lustig et al., 2016) states, with distinct activity in the theta range (3-12Hz); although these studies did not examine connectivity within the hippocampal recording area at the frequencies we investigated.

The three hippocampal LFP states differ with respect to the associated multi-unit activity. Our quantitative analysis of multi-unit parameters (**Table 1**) corroborated the impression based on visual inspection of hippocampal recordings (**Figure 1**) that, overall, neuronal firing, in particular burst firing, is suppressed in the suppression as compared to the burst and continuous states. More specifically, the percentage of spikes fired in burst was significantly higher and overall firing rate was, or strongly tended to be, higher, respectively, in the burst and continuous state compared to the suppression state. It should be noted that, although these differences were suggested by visual inspection of most recordings, they were not always apparent even in recordings from different electrodes in the same rat. For example, in **Figure 1** burst and suppression states clearly show higher MUA on channel 1 whereas on channel 2, which shows a higher overall firing rate, the states display similar MUA, possibly reflecting a ceiling effect. Increased neuronal firing in the burst compared to the suppression state has been reported previously in neocortical regions (Ferron, Kroeger, Chever, & Amzica, 2009; Kroeger & Amzica, 2007; Steriade, Amzica, & Contreras, 1994) and in the subiculum (Land et al., 2012). To explain this difference, it was proposed that anaesthesia induces a hyperexcitable state with an increased extracellular Ca2+ concentration. During the burst state the high Ca2+ would facilitate synaptic transmission causing the Ca2+ to move into the neuron, which would then lead to the suppression state, a refractory period during which Ca2+ concentration is relatively high within (and relatively low outside) of the neurons, therefore decreasing the likelihood of neuronal activity until the Ca2+ is transported back into the extracellular space (Ferron et al., 2009; Kroeger & Amzica, 2007). Taken together, our semi-automated separation method based on the kurtosis of the LFP amplitude distribution, revealed three distinct hippocampal LFP states – burst, suppression and continuous state – whose properties correspond to those of similar hippocampal LFP states identified previously based on visual inspection or other methods (**Table 2**). The cumulative duration of each of the three hippocampal LFP states was not affected by local infusion of the GABA-A receptor antagonist picrotoxin, indicating that the expression of the states does not depend on GABA-A receptor-mediated inhibition in the hippocampus. However, as discussed in the next section, hippocampal neural disinhibition by picrotoxin changed the LFP properties within the three hippocampal LFP states.

**Table 2.** **The continuous and the burst-suppression states of the hippocampal LFP in isoflurane-anaesthetized rats: comparison to previously described hippocampal and cortical LFP states under anaesthesia** The direct quotations highlight state properties corresponding to the ‘continuous’ and ‘burst-suppression’ states shown in the present study. Additional comments regarding the authors’ interpretation of these states, and any comments on the relation to the states described in the present studies, are shown in the additional comments column.

| Paper | 'Continuous' State | 'Burst-Suppression' State | Additional Comments |
| --- | --- | --- | --- |
| Clement et al. 2008 | **State referred to as:** Activated State<br>**LFP pattern:** 'Activated EEG patterns consisted of low amplitude fast activity at neocortical sites concomitant with theta (rhythmic 3–5 Hz) activity at hippocampal sites'<br>**Unit activity:** Not Reported | No equivalent state reported | **Species:** Rats<br>**Anaesthesia:** Urethane<br>**Recording area:** Hippocampus<br>A hippocampal slow oscillation state (see Wolansky et al. 2006, from the same research group) is mentioned. |
| Ferron et al. 2009 | **State referred to as:** Low-Isoflurane<br>**LFP pattern:** [EEG] 'All recording sessions started under 1.5% isoflurane anesthesia, which produced activity patterns comparable to those recorded during slow-wave sleep (SWS) and under ketamine-xylazine anesthesia (Fig. 1A) ... Briefly, these patterns consist of a quasi regular slow oscillation with a frequency range just below 1 Hz.'<br>**Unit activity:** Not Reported | **State referred to as:** Burst-Suppression<br>**LFP pattern:** [EEG] '[A] deep level of anesthesia, producing burst suppression (BS), generates a state of unstable cortical hyperexcitability... BS is an electroencephalographic (EEG) pattern consisting of an alternating pattern of (1) epochs with silenced cortical activity and associated EEG isoelectric line, and (2) periods displaying high-amplitude slow and sharp waves'<br>**MUA:** [Intracellular Recordings] 'The EEG patterns during BS consisted of periods of bursting and isoelectric lines. During the latter, all recorded cortical neurons displayed a hyperpolarized membrane potential devoid of any synaptic activity, whereas bursts consisted of depolarizing triangular-shaped potentials, synchronous with corresponding field and EEG waves.' | **Species:** Cats<br>**Anaesthesia:** Isoflurane<br>**Recording area:** Neocortex<br>It is unclear how this SWS mentioned here relates to our Activated state, as the isoflurane level (1.5%) is higher than we, and others, report for similar states. It may be a state which is part of the transition from our Activated to Deactivated State. |
| Kroeger and Amzica 2007 | **State referred to as:** Low-Isoflurane/SWS<br>**LFP pattern:** [EEG] 'Lower doses (1–2% isoflurane), sufficient for painless surgical procedures, produced a pattern of continuous sleep-like slow waves'<br>**Unit activity:** Not Reported (in absence of stimulation) | **State referred to as:** Burst-Suppression<br>**LFP pattern:** [EEG] '[A] state of deep anesthesia induced by various anesthetics (isoflurane, barbiturates, and propofol) associated with an electroencephalographic pattern of BS... BS was obtained with higher doses of anesthetic (in the case of isoflurane, 2%), and its patterns, in accordance with previous descriptions, were characterized at the EEG level by alternating periods of (1) bursts of high-amplitude and low-frequency (15 Hz) waves, and (2) isoelectric EEG.'<br>**MUA:** [Intracellular Recordings] 'This further supports the claim that the state of BS is a state of hyperexcitability. The onset of this state window is prefigured by a progressive increase in the extracellular Ca<sup>2+</sup> concentrations that may, on the one hand, transiently enhance cortical synaptic processes, and, on the other hand, reduce spontaneous<br>firing of neurons through enhanced screening of Na<sup>+</sup> channels... thus contributing to the gradual onset of suppression episodes.' | **Species:** Cats<br>**Anaesthesia:** Isoflurane<br>**Recording area:** Neocortex<br>See comments above for Ferron et al. 2009 relating to the low-isoflurane state. |
| Land et al. 2012 | **State referred to as:** Light Anaesthesia/Low Suppression<br>**LFP pattern:** 'After the decrease of isoflurane concentration to 0.7 vol ... activity to finally return into the initial state of small amplitude LFP activity.'<br>'The light anesthesia state was characterized by theta oscillations in the subiculum with a mean peak frequency of 4.8 Hz (SD 0.3 Hz)'<br>**MUA:** 'After the decrease of isoflurane concentration to 0.7 vol %, this sequence of events appeared in reversed order: suppression was followed by burst-suppression, then slow rhythmic activity to finally return into the initial state of small amplitude LFP activity... Multiunit activity in V1 and the subiculum followed these changes in LFP activity [in this order or in reverse] with analogous changes in spiking patterns.' | **State referred to as:** Deep Anaesthesia/High Suppression/Burst-suppression<br>**LFP pattern:** 'Increasing levels of anesthesia eventually produce burst suppression of cortical activity, a pattern of alternating high amplitude bursts and periods of suppressed activity (silent periods) signifying deep anesthesia'<br>'In anesthesia-induced burst-suppression, the duration of silent periods between bursts is dose-dependent'<br>'The deep anesthesia state was characterized by regular slow rhythmic burst-suppression in the subiculum, with a mean peak frequency of 0.17 Hz (SD 0.05 Hz)'<br>**MUA:** 'LFP bursts in V1 or the subiculum were always accompanied with bursts of strong multiunit activity at the respective electrode sites' | **Species:** Mice<br>**Anaesthesia:** Isoflurane<br>**Recording area:** Hippocampus |
| Lustig et al. 2016 | **State referred to as:** Light isoflurane state<br>**LFP pattern:** 'Recordings during the lighter level of isoflurane anesthesia (~1%) consistently displayed a slow oscillation with a peak frequency of 3.4 Hz +/-0.57 Hz'<br>'The depth profile of the running associated theta oscillations was strikingly similar to the slow oscillation during the light isoflurane state (3.5 Hz).'<br>'[D]uring the light isoflurane state, we observed a slow, theta-like oscillation in CA1 and a faster, gamma-like oscillation around DG.'<br>**Single unit activity:** '[A]ctivity of neurons recorded in the CA1 and DG layers was modulated by theta-like and gamma-like oscillations, respectively' | **State referred to as:** Deep isoflurane state<br>**LFP pattern:** 'under a deeper level of isoflurane anesthesia (2%), hippocampal activity was dominated by an oscillation with a peak frequency of 11.18 Hz ± 0.1 Hz'<br>'[T]he depth profile of SPWs [sharp wave ripples] during water reward and quiet ... was strikingly similar to the individual cycles of the 11 Hz oscillation during the deep isoflurane state.'<br>'In addition to these fast oscillatory events, we frequently observed randomly interspersed sharp LFP spikes in the D3 region.'<br>**Single unit activity:** 'Firing in both CA1 and CA3 neurons was strongly phase locked to the individual cycles (Fig. 4C, top). CA1 neurons tended to fire a burst of action potentials around the trough of each cycle' | **Species:** Rats<br>**Anaesthesia:** Isoflurane<br>**Recording area:** Hippocampus<br>'We found that under different levels of isoflurane anesthesia activity in the hippocampus of rats displays two distinct states, which have several qualities that mirror the theta and LIA states.' |
| Pagliardini et al. 2013 | **State referred to as:** Activated or REM-like<br>**LFP pattern:** 'The REM-like state was characterized by high power in the theta frequency bandwidth (range: 3.5-4.5 Hz) with a corresponding increase in power through the gamma band (25-40Hz), consistent with the expression of gamma activity during both natural sleep and during anesthesia in previous studies'<br>**Unit activity:** Not Reported | **State referred to as:** Deactivated or SWS-like<br>**LFP pattern:** 'The SWS-like state was characterized by high power slow oscillation (SO) bandwidth (range: 0.2-1.2 Hz).'<br>**Unit activity:** Not Reported | **Species:** Mice<br>**Anaesthesia:** Urethane<br>**Recording area:** Hippocampus<br>'[S]upporting the hypothesis that the alternations are intrinsic to the anesthetic effect induced by urethane and not generated by a gradual waning of the anesthetic level.' |
| Steriade et al.<br>1994 | **No equivalent state reported** | **State referred to as:** Burst-Suppression<br>**LFP pattern:** 'The EEG pattern of burst-suppression that followed administration of various anesthetics generally consisted of sharp waves with high amplitudes, at frequencies usually ranging between 2 and 7 Hz, separated by progressively longer periods of complete flatness. The electrical silence lasted for 1-2 sec during the initial stage of this pattern (Fig. 1A3), but reached 5-15 sec in later stages (Figs. 1A 4 and 2C) and even 30 sec or more in some instances (see below, Figs. 7 and 9A).'<br>**Unit activity:** [Extra- and Intra-cellular recordings] 'Only 60-70% of them (11 RE thalamic and 20 thalamocortical cells) ceased firing before the occurrence of burst-suppression; thereafter, those cells discharged during EEG wave bursts and were completely silent during flat periods of the EEG.' | **Species:** Cats<br>**Anaesthesia:** Various<br>**Recording area:** Neocortex and Thalamus |
| Wolansky et al. 2006 | **State referred to as:** Activated State or Theta State<br>**LFP pattern:** 'The activated state [during sleep] consists of prominent 3–12 Hz rhythmical activity known as theta or rhythmic slow activity.' '[T]he same electrographic could be observed during spontaneous state changes under urethane anesthesia'. 'Under urethane, [Slow Oscillation state, see Additional comments] ... appeared to be dependent on the level of anesthesia; increasing anesthetic dosage moderately increased the proportion of time spontaneously spent in the slow oscillatory state (correspondingly decreasing the amount of time spontaneously spent in theta).'<br>**Single unit activity:** 'Most of the neurons that generated rhythmic action potentials during the SO also did so during theta ' | **No equivalent state reported** | **Species:** Rats<br>**Anaesthesia:** Urethane<br>**Recording area:** Hippocampus<br>Wolansky et al. report a Hippocampal Slow Oscillation state at 1Hz with a strong gamma power. It is unclear how this relates to our states, but it is not directly comparable to our burst-suppression state. |

### 4.2. State- and frequency-dependent changes in the hippocampal LFP power by local picrotoxin disinhibition

In both burst and suppression states picrotoxin, as compared to saline infusions, tended to increase hippocampal LFP power at lower frequencies (<20 Hz) and to decrease power at higher frequencies (>20 Hz) although only the increase at 6-18 Hz in the burst state reached statistical significance. In contrast, there was no evidence of picrotoxin-induced power changes in the continuous LFP state. Our findings are consistent with several previous in vivo and in vitro studies reporting changes in the hippocampal LFP following manipulations of GABAergic inhibition. Our finding that hippocampal GABA antagonism by picrotoxin facilitates LFP oscillations at lower frequencies, including the theta range, is consistent with many previous in vitro and vivo findings indicating an inverse relationship between GABAergic inhibition and theta oscillations in the hippocampus; which is consistent with the idea that hippocampal disinhibition by the medial septum, from where long-range GABAergic projections synapse onto parvalbumin-positive GABAergic inhibitory hippocampal interneurons (Freund & Antal, 1988), is a key extrinsic mechanism of hippocampal theta generation (Buzsaki, 2002; Colgin, 2016; Kowalczyk et al., 2013)see Introduction). However, some aspects of hippocampal theta and other slow oscillations may be facilitated by GABAergic inhibition. In rat hippocampal slices, GABA-A antagonists decreased the amplitude of electrically induced theta oscillations (Heynen et al., 1993), and abolished ‘regular ventral hippocampus spontaneous synchronous activity’, a 1-2 Hz oscillation (Papatheodoropoulos & Kostopoulos, 2002), possibly reflecting specific properties of these in vitro hippocampal slow oscillations. Moreover, optogenetic activation of parvalbumin positive GABAergic hippocampal interneurons strengthened, whereas silencing of these neurons disrupted, theta oscillations in an in vitro hippocampal preparation with intact intrinsic but severed extrinsic connectivity, showing that GABAergic inhibition by parvalbumin interneurons supports intrinsically generated hippocampal theta (Amilhon et al., 2015).

Although we did not find any statistically significant effects of picrotoxin on hippocampal LFP power in the beta/gamma frequencies (specifically 20-30Hz), picrotoxin tended to decrease power in this frequency band within the burst and suppression states. These numerical decreases are consistent with previous findings that pharmacological disinhibition disrupts hippocampal in vitro gamma oscillations and that GABAergic inhibition within the hippocampus plays a key role in generating these oscillations (Bartos et al., 2007; Buzsaki & Wang, 2012; Traub et al., 2004; Whittington et al., 2000). There is a wealth of in vitro studies showing that the GABA-A antagonist bicuculline applied to hippocampal slices decreases not only the power of 20-30Hz oscillations, but also the frequency of these oscillations (Arai & Natsume, 2006; Boddeke, Best, & Boeijinga, 1997; Shimono, Brucher, Granger, Lynch, & Taketani, 2000; Trevino, Vivar, & Gutierrez, 2007).

### 4.3. Reduced functional connectivity within the temporal to intermediate hippocampus after picrotoxin disinhibition

In the suppression and burst states, functional connectivity within the temporal to intermediate hippocampus, as reflected by phase locking of the LFP signal across the different electrodes of our recording array, was decreased in the gamma range (> 20 Hz), this decrease was statistically significant in the burst state from 29.5 – 33Hz. This is consistent with previous in vitro findings that disinhibition by morphine disrupted synchrony of gamma oscillations across a hippocampal slice preparation (Faulkner, Traub, & Whittington, 1998; Whittington, Traub, Faulkner, Jefferys, & Chettiar, 1998). Synchrony was disrupted between areas that were 1.5-2.5 mm apart, whereas effects were not observed at shorter distances (Whittington et al., 1998). It is therefore possible that, in the present study, this effect was weakened, because we assessed the collective connectivity over our 2-mm electrode array, including short distances between neighbouring electrodes. Faulkner and colleagues (1998) also reported that a GABA-A agonist applied to hippocampal slices causes a similar decrease in gamma-frequency synchrony, suggesting that such synchrony depends on a balanced level of hippocampal GABA inhibition. Furthermore, neuro-computational studies suggest that, through multiple mechanisms, GABAergic interneurons are important for the synchronisation of gamma oscillations (Bartos et al., 2007; Whittington et al., 2000). For example, in a neuro-computational model of hippocampal pyramidal cells and GABAergic interneurons, blocking GABA-A receptors on the pyramidal cells abolished coherence at gamma frequencies (Traub et al., 2000). Our findings are the first in vivo findings to support these in vitro and modelling findings that the disruption of GABAergic inhibition, through blocking of hippocampal GABA-A receptors, disrupts synchronization of gamma oscillations across the hippocampus.

Finally, we found a numerical trend for picrotoxin to reduce functional connectivity in the theta frequency in the continuous hippocampal LFP state. This is consistent with previous findings that blockade of GABA-A receptors decreases enthorinal-subicular theta coherence in vitro (Levesque et al., 2017) and that genetic knock-out of the GABA-A receptor subunit β3 decreased theta cross-correlation between different hippocampal subfields in freely moving mice (Hentschke et al., 2009).

### 4.4. State-dependence of enhanced multi-unit burst firing caused by hippocampal neural disinhibition

Similar to the state-dependence of LFP changes, the picrotoxin-induced enhancement of hippocampal burst firing (McGarrity et al., 2017) tended to be more pronounced in the burst and suppression states than in the continuous state, although only the analysis of burst duration revealed a statistically significant interaction of the LFP state with the infusion effect. Previous findings suggest that in an isoflurane-induced burst-suppression state, the cortex is in a hyperexcitable state due to a decrease of excitation leading to a decrease in inhibition, which overall results in the excitation/inhibition balance being shifted towards excitation (Ferron et al., 2009). Neuronal hyperexcitability is also supported by the finding that, during the burst-suppression state, subliminal sensory stimuli activate cortical areas, including the subiculum, that are not activated by the same stimuli during other anaesthetized or the awake states (Kroeger & Amzica, 2007; Land et al., 2012). The more pronounced effects of picrotoxin during the burst and suppression states, compared to the continuous state, may reflect neuronal hyperexcitability during these states.

### 4.5. Conclusions

We have presented an objective, semi-automated method for separating three distinct hippocampal LFP states in isoflurane-anaesthetized rats, including the burst, suppression and continuous states. These states are characterized by different LFP properties and associated multi-unit activity, which are consistent with the properties of burst-suppression and ‘activated’ or ‘light-anaesthesia’ hippocampal LFP states that have previously been identified based on visual inspection and other methods (**Table 2**). Furthermore, our finding that the enhanced hippocampal multi-unit burst firing induced by hippocampal picrotoxin infusion is more pronounced in the burst and suppression state, compared to the continuous state, is consistent with previous studies suggesting that the burst and suppression states are characterized by neuronal hyperexcitability (Ferron et al., 2009; Kroeger & Amzica, 2007; Land et al., 2012). Our state-separated analysis of the impact of hippocampal picrotoxin infusion on LFP properties around the infusion site revealed that, in the burst and suppression states, neural disinhibition tended to increase low frequency oscillations (<20 Hz), including theta oscillations, and to decrease gamma frequency oscillations (>20 Hz), although only the picrotoxin-induced power increase between 6 and 18 Hz in the burst state was statistically significant. In addition, neural disinhibition reduced functional connectivity, as reflected by phase-lock Eigenvalues, across the recording area in the temporal to intermediate hippocampus at gamma frequencies in the burst state (with significant reductions compared to saline infusion between 29.5 and 33Hz). These findings support that GABA-A receptor-mediated mechanisms regulate hippocampal LFP oscillations in vivo (albeit under anaesthesia), confirming and extending previous findings mainly from in vitro studies that GABA-A-mediated inhibition negatively modulates lower frequencies, including theta frequencies (Kowalczyk et al., 2013), and positively modulates power and connectivity at higher frequencies (Bartos et al., 2007; Buzsaki & Wang, 2012; Traub et al., 2004; Whittington et al., 2000). Cortical, including hippocampal, LFP oscillations have been suggested to be important for memory and other cognitive functions, and alterations of such oscillations have been linked to memory and other cognitive impairments (Buzsáki & Draguhn, 2004; Colgin, 2016; Uhlhaas & Singer, 2006, 2010). Consistent with this prominent view, the frequency-specific alterations of hippocampal LFP properties caused by hippocampal neural disinhibition may contribute to the memory and attentional impairments caused by the same manipulation (McGarrity et al., 2017). However, direct evidence causally linking disinhibition-induced hippocampal LFP changes to the memory and attentional deficits is lacking and potential underlying mechanisms would also require clarification. One important question raised by the present findings is i) if similar changes in LFP oscillations would be observed without anaesthesia and ii) how such changes relate to alterations in surface EEG recordings that have been reported in relevant clinical conditions, including schizophrenia and age-related cognitive decline (Hunt, Kopell, Traub, & Whittington, 2017; Uhlhaas & Singer, 2006, 2010). To address these two questions, our ongoing studies examine the simultaneous impact of hippocampal neural disinhibition on the hippocampal LFP and on surface EEG recordings in freely moving rats.

## Acknowledgements

This work was supported by a BBSRC iCASE PhD award in partnership with Boehringer Ingelheim to Miriam Gwilt and Tobias Bast. During preparation of this paper, Tobias Bast was a Fellow of the Research Group ‘Cognitive Behavior of Humans, Animals, and Machines: Situation Model Perspectives’ at the Center for Interdisciplinary Research (ZiF) at Bielefeld University and benefitted from a research sabbatical granted by the School of Psychology at the University of Nottingham. Markus Bauer was supported by Nottingham Research Fellowship by the University of Nottingham.

